# PepGM: A probabilistic graphical model for taxonomic inference of viral proteome samples with associated confidence scores

**DOI:** 10.1101/2022.09.21.508832

**Authors:** Tanja Holstein, Franziska Kistner, Lennart Martens, Thilo Muth

## Abstract

**Motivation:** Inferring taxonomy in mass spectrometry-based shotgun proteomics is a complex task. In multi-species or viral samples of unknown taxonomic origin, the presence of proteins and corresponding taxa must be inferred from a list of identified peptides which is often complicated by protein homology: many proteins do not only share peptides within a taxon but also between taxa. However, correct taxonomic identification is crucial when identifying different viral strains with high sequence homology – considering, e.g., the different epidemiological characteristics of the various strains of SARS-CoV-2. Additionally, many viruses mutate frequently, further complicating the correct assignment of virus proteomic samples.

**Results:** We present PepGM, a probabilistic graphical for the taxonomic assignment of virus proteomic samples with strain-level resolution and associated confidence scores. PepGM combines the results of a standard proteomic database search algorithm with belief propagation to calculate the marginal distributions, and thus confidence score, for potential taxonomic assignments. We demonstrate the performance of PepGM using several publicly available virus proteomic datasets, showing its strain-level resolution performance. In two out of eight cases, the taxonomic assignments were only correct on species level, which PepGM clearly indicates by lower confidence scores.

**Availability and Implementation:** PepGM is written in Python and embedded into a Snakemake workflow. Its is available at https://github.com/BAMeScience/PepGM

## Introduction

Viruses and especially viral pathogens represent a tremendous threat to public health. While the threat of emerging viral diseases has been known to the scientific community for the past several decades,^1–3^ the SARS-CoV-2 pandemic has propelled the importance of viral surveillance and investigation to the forefront of public attention.^4,5^ In this context, an essential element is the fast and accurate taxonomic identification of viral samples. As different strains of the same viral species can differ strongly regarding patient outcome and epidemiological characteristics,^4,6–8^ correct strain-level attribution is crucial. While the gold standard method for taxonomic identification of viral samples is based on genomic analysis by next generation sequencing,^9,10^ bottom-up proteomics-based analysis is a promising orthogonal approach. ^11^ The public need for alternative bioanalytical tools is stressed by reagent shortages during global pandemic situations due to only few available analysis methods.^12,13^ The correct strain-level taxonomic inference of proteomic samples, however, remains a challenging task. An experimental approach that has been pursued in the past is the use of MALDI-TOF for so-called biotyping or proteotyping.^14–16^ These approaches rely on signature peptides which are detected by mass spectrometry and uniquely associated to a species. Yet these approaches are limited to a single MS level, and tend to be inappropriate for strain-level analysis, as tailored spectral databases with strain-level resolution are required.^16^ Thus, approaches based on tandem mass spectrometry (MS/MS), that offer a more precise view of the actual peptide sequence, have become a promising alternative.^17^ For instance, an MS/MS-based approach for the identification of SARS-CoV-2 from patient samples was recently developed,^18^ based on a targeted proteomics workflow.

For untargeted identification of viral samples based on MS2 spectra only few bioinformatic workflows exist^19–22^ and these focus on the analysis of samples with known organism source.^23^ For the identification of virus strains of unknown taxonomic origin, specific challenges emerge. Many of these overlap with challenges in metaproteomics, a field that aims to identify and quantify the phenotype of microbial communities with unknown taxonomic composition.^24^

First, the analysis of samples with unknown composition potentially requires large search spaces. This is because standard proteomics workflows rely on search algorithms that aim to match experimental MS/MS spectra against a reference database.^25^ Thus, all candidate sequences have to be included in the reference database, which is computationally expensive and increases the risk of false positives. ^26^

Second, unique protein inference is not guaranteed. A single peptide can map to several proteins, impeding accurate protein inference and leading to the so-called protein inference problem.^27,28^ Recently, graphical models have been applied to address this.^29^ Third, taxonomic inference has an analogous issue to protein inference: peptides are shared not only between proteins, but also between taxa, impeding accurate taxonomic inference.

Specific bioinformatic workflows developed to address some of the described issues employ different strategies with varying success rates. Originally developed for metaproteomics, Pi-pasic relies on abundance similarity correction to taxonomically resolve samples with multiple closely-related strains.^22^ Similarly, MiCId uses peptidome similarity correction^20^ to achieve species-level resolution and provide a statistical estimate of its taxonomic assignments, however, it lacks strain level resolution and has difficulties processing very large databases.^19^ Other tools originally developed for metaproteomics, such as Unipept,^21^ map peptides to their lowest common ancestor - meaning the lowest taxonomic level they are specific to. This results in taxonomic identifications without confidence estimates and is inappropriate when strain-level resolution is required, as can be the case in clinical settings, where disease severity and therapeutic decisions strongly depend on strain information. A different approach, TaxIt, ^19^reduces the search space while still taking into account as many reference proteomes as possible through an iterative approach that uses multiple identification steps. ^19^ Amongst all mentioned workflows, it is the most apt at achieving strain-level resolution, but lacks confidence estimates for its taxonomic assignments.

We here present PepGM, a graphical model-based bioinformatic pipeline for taxonomic inference of viral proteome samples. Our approach results in the accurate detection of viral pathogens in proteome samples with strain-level resolution. In particular, PepGM scores and ranks taxonomic assignments, thereby providing statistical confidence estimates. This becomes particularly relevant in situations of ambiguous or erroneous identifications, e.g., when the correct viral strain is not present in the database or when the available strain reference proteomes overlap heavily. PepGM starts with high confidence peptide identifications of a standard database search against a generic viral reference database. Based on the resulting matches, all candidate taxa on strain level are inferred using weighted peptide-spectrum matches. For these taxa, each available strain-level proteome is retrieved. These candidate peptide sequences and the taxa themselves are represented as nodes in a probabilistic graphical model. ^30^ An edge between a taxon and a peptide is drawn if the respective candidate peptide is validated by the database search results. Using the loopy belief propagation algorithm i.e., an algorithm for approximate Bayesian inference which has been successfully applied for protein inference, ^29^ PepGM computes the marginal distribution of taxa, which corresponds to the probability of their presence in the analyzed sample. We demonstrate that PepGM consistently identifies the correct viral strains on an exemplary set of viral proteomic samples from publicly available data, investigating various use cases. We show that the taxon posterior probabilities represent suitable confidence estimates of our newly developed method. The combination of strain level resolution with statistically assigned confidence estimates makes PepGM particularly useful for potential applications in the clinical context, where therapeutic decisions require both very detailed and very confident identifications of pathogens.

## Methods

In the following section, we describe PepGM in detail. We start with an overview of the PepGM workflow, then define the individual workflow steps, input and outputs, and provide a brief description of the graphical model and inference algorithm used.

### Workflow overview

PepGM takes as input the MS/MS spectrum files (.mgf format), a reference database containing one or multiple proteomes, and, optionally, a cRAP (containing common contaminants) and a host database. PepGM broadly comprises the following seven steps (Figure 1) : (0) optional host filtering, (1) standard proteomic database search, (2) identification of candidate taxa, (3) construction of the bipartite peptide-taxon graph, (4) construction of the graphical model, (5) inference algorithm embedded into a grid search, (6) parameter evaluation and finally, results output and visualization.

**Figure 1:**
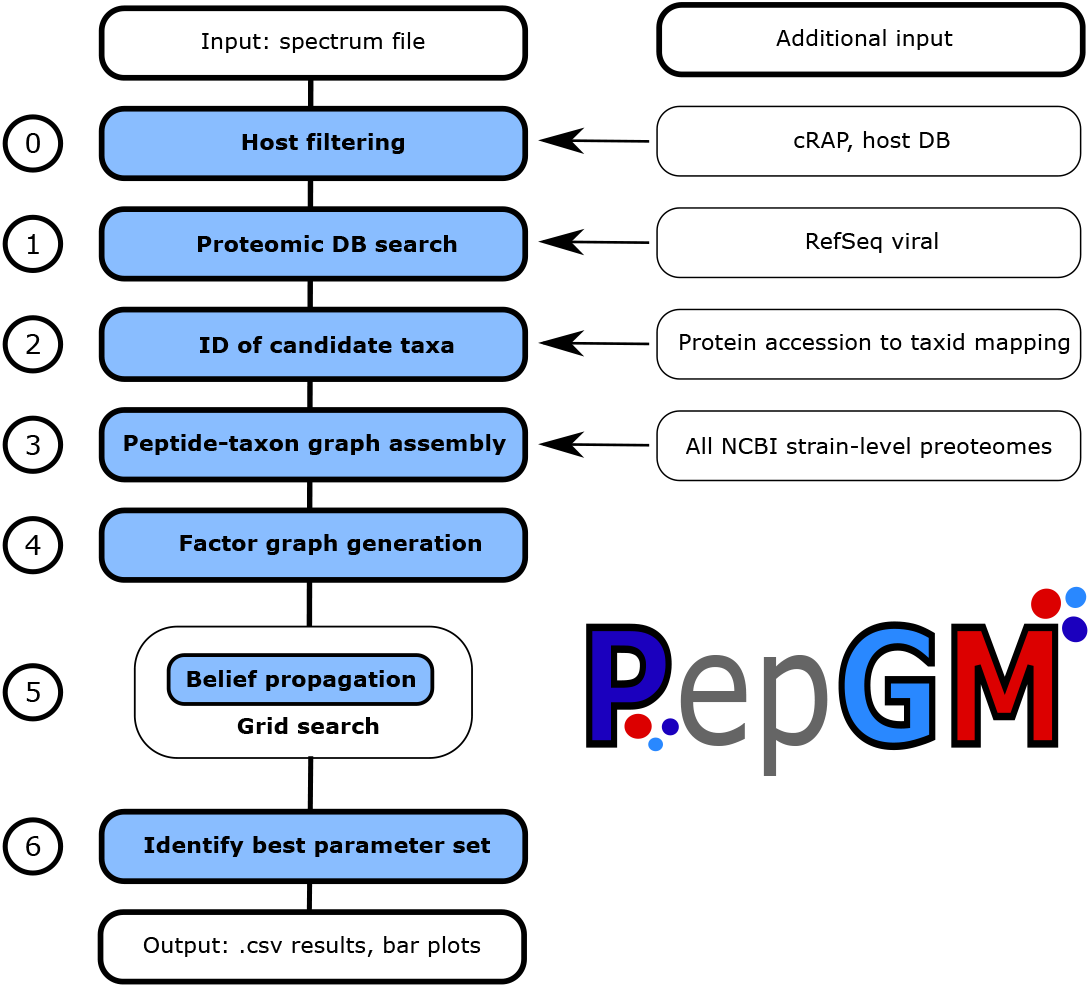
PepGM workflow for taxonomic identification of viral samples. The PepGM workflow comprises 7 steps (numbered 0 to 6). The input is a MS/MS spectrum file (.mgf), additional input or information used at individual workflow steps are specified to the right of the workflow diagram.

We use Snakemake^31^ as a workflow management system, while all individual scripts are written in Python. Additional packages used can be found in supplementary materials Section S1. We will now describe the individual workflow steps.

#### 0- Prefiltering (optional)

If a host and/or contaminant (e.g. cRAP) database is provided by the user, the MS/MS spectra are searched against these. The resulting matches are known to be of non-viral origin and can therefore be excluded from the input.

#### 1- Database search against a general viral reference database

Experimental spectra are searched against a complete virus-specific reference database. This is any database provided by the user in .fasta format. However, due to the taxid-mapping in the subsequent workflow step, proteins need to be provided with NCBI accession numbers in the .fasta file. A taxid is a unique identifier attributed to each taxon by NCBI, a protein accession is the unique identifier for the protein. In this work, we use the Ref-Seq Viral database as a reference.^32^ PepGM uses SearchGUI^33^ for database-driven peptide identification and PeptideShaker^34^ as post-processing tool. Due to its flexible Snakemake implementation, any combination of database search and rescoring algorithm can be used. This step results in a list of scored peptides with corresponding peptide-spectrum matches (PSMs).

#### 2- Inferring candidate taxa

Based on these PSMs, we infer all candidate taxa and only keep high-scoring taxa as assessed by a basic scoring scheme. Taxon confidence is assessed based on weighted PSMs per protein and taxon. PSMs weights are first aggregated at protein level, where the weight of a PSM is adjusted by dividing it by the number of times it occurs in other proteins. The aggregated weights are then analogously propagated to the all potential taxa. Those taxa with a score below the median score are excluded from the candidate set.

#### 3- Construction of the bipartite peptide taxon graph

The identified candidate taxa will be taxonomic nodes at species level, as this is the resolution provided by the NCBI taxon-protein mapping accessions. In the next step, from these candidate species, all corresponding strain-level taxa are inferred and are kept as candidate strains to be included in the graphical model. Each candidate strain proteome is automatically downloaded through the NCBI Entrez API and is in silico digested using tryptic cleavage settings. Next, we assemble a bipartite graph of peptide and taxon nodes, where an edge is drawn between a taxon node and a peptide node if the peptide is part of an in silico trypsin digested taxon proteome. The digested strain-level peptides are matched (using exact string matching) against the list of peptides identified in the previous database search, and each matched peptide is included in the bipartite graph. Finally, the PSM scores are aggregated at peptide level, with only the highest score kept for each peptide.

#### 4- Construction of the factor graph

The subsequent Bayesian network representation we choose for the joint distribution of peptides and taxa corresponds to a representation successfully used for protein inference in previous work.^29^ The bipartite peptide-taxon graph represents the conditional dependencies between peptides and their ‘parent’ taxa. Consequently, the high-dimensional joint distribution of peptides and taxa is factorized into less complex distributions, which are the prior distributions of the taxa and conditional probability distributions (CPD) for the peptides given the presence of a parent taxon. Both peptides and taxa are binary variables. As the actual conditional probability distribution of the peptides depending on the presence of the taxa is not known, it is modelled using the noisy-OR model^35^ . As for the protein inference model, this means that we assume that the presence of any of the parent taxa is sufficient to produce a peptide belonging to a taxon proteome. Three parameters, that we assume to be the same for all peptides and taxa, results from this representation:

- *α* : probability for a peptide to be observed given the presence of its parent taxon
- *β* : probability for a peptide to be randomly (falsely) observed
- *γ* : prior probability for a taxon to be present

Albeit the assumption that *α, β* and *γ* are the same for all peptides is a strong simplification, it has already proven itself useful for protein inference.^29^ Following the described model, a CPD is computed for each peptide. Additional convolution tree^36^ nodes speed up the propagation of probabilities between peptide and taxon level during the ensuing inference algorithm.

#### 5- Inference algorithm and grid search

Using the loopy belief propagation algorithm,^37^ posterior probabilities are calculated for all taxa in the graph. The loopy belief propagation is embedded in a grid search through the parameter space of *α, β* and *γ*. Each parameter *α, β, γ* ∈ [0, 1]. The grid search covers a set of 126 parameter combinations with individual parameters logarithmically spaced. Each parameter set provides a list of potential taxonomic identification results.

#### 6- Parameter evaluation and results output

Through an empirically determined metric, based on the distribution of taxon scores, we identify the parameter range best fitting for the sample at hand. The metric to be maximized is the following:

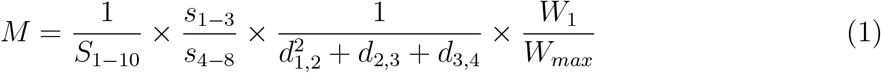

In equation 1, the subscripts have the following meanings. *S*_*i*–*j*_ is the entropy of the distribution of the *i*-st to *j*-th posterior taxon probabilities. Icluding the entropy ensures that the results distribution selected contains as much information as possible. *s*_*i*–*j*_ is the sum of the *i*-th to *j*-th posterior taxon probabilities, favoring distributions where the higher scoring taxa clearly delimit themselves from the lower scoring ones. *d_i;j_* is the taxonomic distance (calculated as the number of taxonomic nodes between two taxa) between the *i*-th and *j*-th taxa, the corresponding term favors distribution where the top scoring taxa are taxonomically close. Finally, *W_i_* is the weight attributed to taxon *i* (or the parent species of taxon *i*) during the acquisition of candidate taxa, while *W_max_* is the maximum weight any taxon was attributed during analysis. This ensures that the top scoring taxon identified by PepGM is also ones that was attributed many PSMs.

The taxonomic identification results with their associated confidence scores are given in the output as bar plots, as a .csv file and as a figure of their visual projection onto a phylogenetic tree.

### Selection of samples for evaluation and further parameter settings

To evaluate PepGM, we selected several pathogenic viral samples that were publicly available in the PRIDE repository. ^38^ We selected the following samples: two *Cowpox virus (strain Brighton Red)* (PXD014913 and PXD003013), one *Human adenovirus 2* (PXD004095), two *SARS-CoV-2* (PXD024130 and PXD018594), one *Human herpesvirus 1 (strain 17)* (PXD005104), one *Hendra virus (strain Horse/Autralia/Hendra/1994)* (PXD001165) and one avian bronchitis (Beaudette CK) (PXD002936). Throughout the manuscript, we will refer to them as Cowpox virus PXD014923 or PXD003013, adenovirus, SARS-CoV-2 PXD024130 or PXD018594, herpesvirus, hendravirus, and avian bronchitis respectively. The description of the samples acquisition and sample-specific search parameter settings can be found in the supplementary materials. Except for SARS-CoV-2, all sample taxa were available with strain-level resolution in the NCBI taxonomy database. The host background proteomes were downloaded using Uniprot as described in the supplementary materials.

For the proteomic database search with SearchGUI, we used the RefSeq Viral reference database, downloaded through NCBI. As search engine, we selected X!Tandem. The FDR was set to 5%, which is high compared to standard proteomic workflows. We performed additional sample analyses at 1% FDR and 20 % FDR (too evaluate the effect of adding many low confidence PSMs), where we show that the influence of using different FDRs was limited.

Except for SARS-CoV-2, all of the selected samples previously served as benchmarking samples for the TaxIt^19^ pipeline, where the taxonomic identifications were benchmarked using MiCiD,^20^ Unipept,^21^ and Pipasic.^22^ It was shown that TaxIt performed best among all software, being the only one to consistently provide strain-level taxonomic identifications, therefore, we compare PepGM directly to TaxIt. Runtime and memory usage are evaluated using a Fujitsu laptop with Ubuntu 20.04.4 LTS, with a 4-core Intel i5-7200 CPU@2.50GHz and 16GB of memory.

## Results

### Strain-level identification accuracy

To demonstrate that PepGM consistently identifies the correct strain in viral samples, we run the complete PepGM workflow without the optional host filtering on all samples described above. For the adenovirus, the posterior scores of the 15 highest scoring taxa are shown in Figure 2. The same representation of the PepGM results for all other samples can be found in the supplementary materials in Section S2. Figure 2 (A) shows that adenovirus strain 2 is correctly identified as the strain present in the sample. Two other human mastadenovirus C strains, adenovirus 5 and adenovirus 6, also receive high posterior scores. As can be seen in the taxonomic tree representation in Figure 2 (B), these lie taxonomically very close to the adenovirus 2 and likely have high sequence similarity. To investigate this further, we computed the peptidome similarity of the 15 top scoring viral strains identified for the adenovirus with a symmetric similarity measure.^39^ Here, the peptidome similarity was restricted to peptides that were detected, and therefore present in the graphical model, to better represent the similarity of the actually measured peptidomes. We call this measure the detected peptidome similarity. For a peptide set X and a second peptide set Y, our similarity measure corresponds to 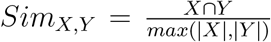. As expected, the three top scoring strains - human adenovirus 2, 5 and 6 - have a high detected peptidome similarity between 0.88 and 0.96 (Figure 3). The two other slightly higher scoring taxa (Simian adenovirus 34 and human adenovirus 57 with posterior probability scores of 0.21 and 0.14, respectively) also have a slightly higher detected peptidome similarity (0.21 and 0.11 respectively) than other, lower scoring taxa. All other taxa included in Figure 2 are not part of the mastadenovirus genus and do not share detected peptides with the mastadenoviruses. Their presence in the results likely stems from either wrongly detected peptides or the presence of host proteins or other impurities. The low assigned posterior probabilities (all below 0.13) indicate that these are correctly identified as absent. An overview of the results showing the correct viral strain, the identified viral strain and the computed posterior score for all analyzed samples is shown in Table 1. PepGM identifies the correct species for all, and the correct strain for six out of 8 analyzed samples and attributes high posterior probabilities (between 0.99 and 0.88) to the accurately identified taxa. For the herpesvirus and the avian bronchitis, we marked the identification results as ambiguous because PepGM was either not able to resolve between closely related strains or identified a neighbouring strain (from the correct species) as present. We next investigate the potential reasons for such ambiguous identifications.

**Figure 2:**
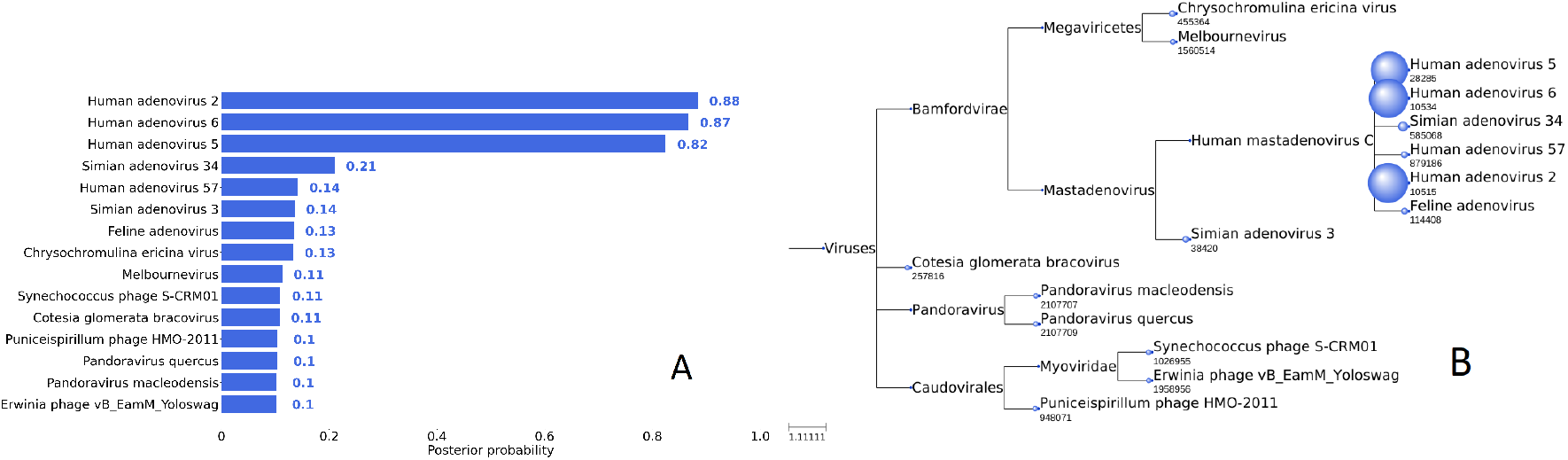
PepGM identification results for the adenovirus sample. Bar plot representation of the 15 highest scoring taxa, with the exact attributed score written next to each bar (A), and results projected onto the taxonomic tree (B).

**Figure 3:**
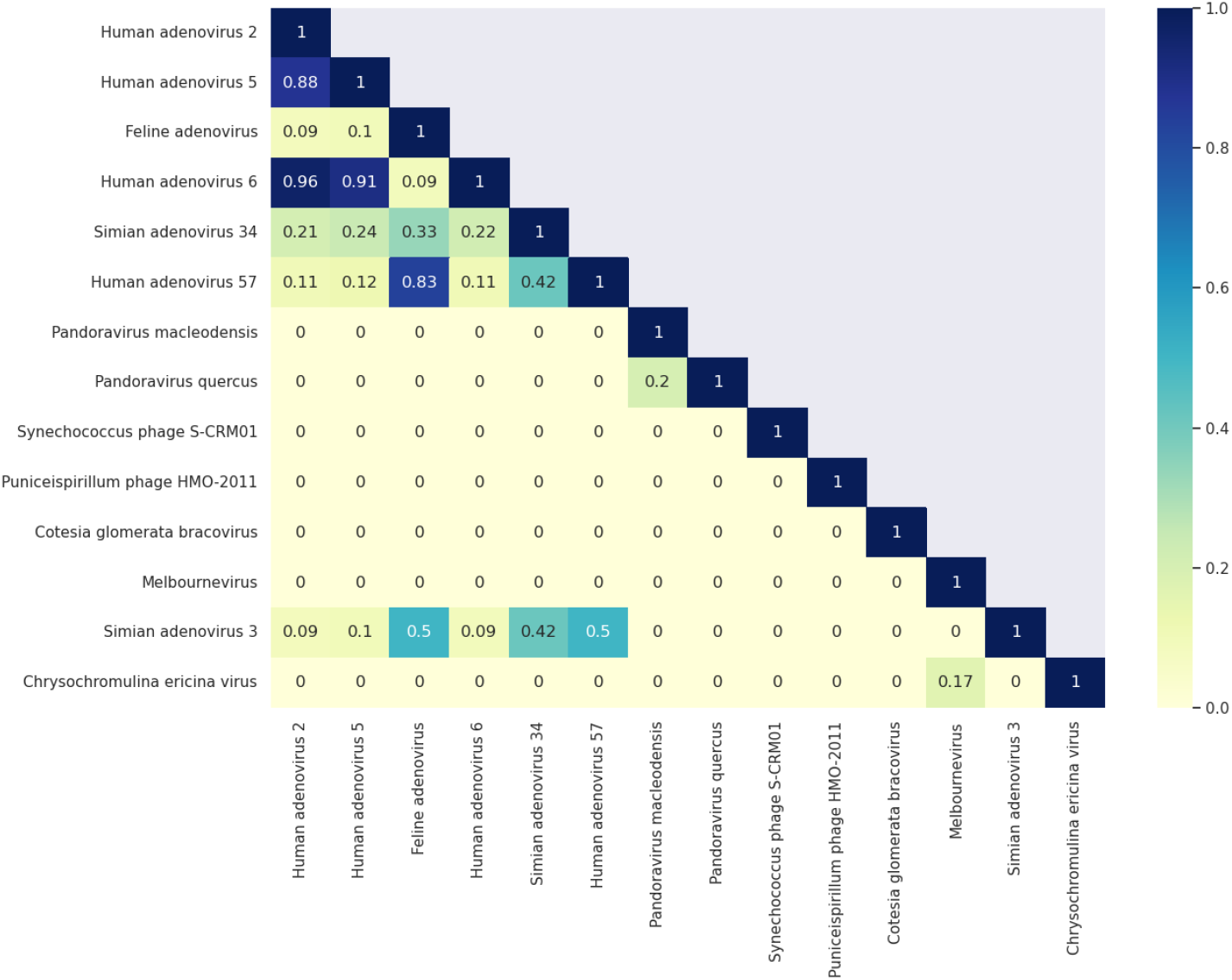
Detected peptidome similarity of the top scoring viral strains for the adenovirus sample. Peptides included for the peptidome similarity computation are restricted to the peptides that were identified by SearchGUI/PeptideShaker and included in the graphical model.

**Table 1:** Overview of the results of taxonomic analysis of 8 selected samples using PepGM. For all samples, PepGM identifies the correct species. For the strain identification, *✓* indicates a correct identification, +/− indicates ambiguous results which included the correct identification.

| Sample | Virus (strain) | Identified correctly? | Posterior probability |
| --- | --- | --- | --- |
| PXD003013 | Cowpox virus (Brighton Red) | ✓ | 0.96 |
| PXD014913 | Cowpox virus (Brighton Red) | ✓ | 0.88 |
| PXD018594 | SARS-CoV-2 | ✓ | 0.99 |
| PXD025130 | SARS-CoV-2 | ✓ | 0.99 |
| PXD005104 | Human herpesvirus 1 (strain F) | +/- | 0.81 |
| PXD002936 | Avian bronchitis (Beaudette CK) | +/- | 0.78 |
| PXD001165 | Hendra virus 1994 | ✓ | 0.99 |
| PXD004095 | Human adenovirus (strain 2) | ✓ | 0.88 |

Figure 4 shows the top scoring taxa and their mapping onto the taxonomic tree for the avian bronchitis. Figure 4 shows that PepGM cannot differentiate between the three avian bronchitis Beaudette (CK, US, and no extension) strains. The US and CK strains are sub-types of the Beaudette strain that have been adapted to grow optimally in different cell lines.^40^ They are therefore taxonomically (and protein sequence-wise) very close, which is why PepGM is not able to differentiate between them. The posterior probability that each of the Beaudette strains is attributed - about 0.78 to 0.79, is lower than the probability that was attributed to correct identifications (which where all above 0.88), demonstrating the accuracy of the confidence estimate based on the posterior probabilities. None of the other taxonomic identification software tools, including TaxIt itself, were able to identify the correct Beaudette strain, therefore PepGM has the added benefit of reporting the lower confidence of its identification. To further explore our results, we compute the detected peptidome similarity for the avian bronchitis sample. The results depicted in Figure 5 show that the three avian bronchitis Beaudette strains have a detected peptidome similarity of 1, which means that all identified peptides map to all three strains. This shows that, based on the available information, no algorithm would be able to differentiate between them.

**Figure 4:**
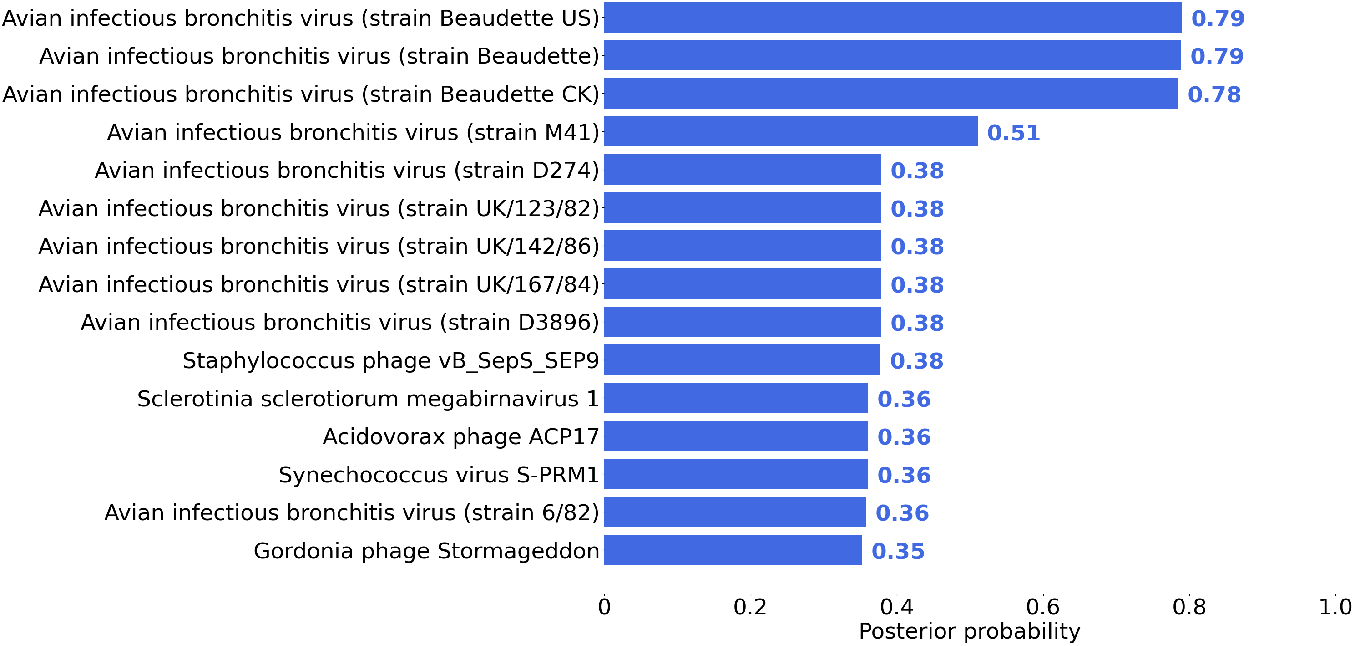
PepGM identification results for the avian bronchitis sample. Bar plot representation of the 15 highest scoring taxa, with the exact attributed score written next to each bar.

**Figure 5:**
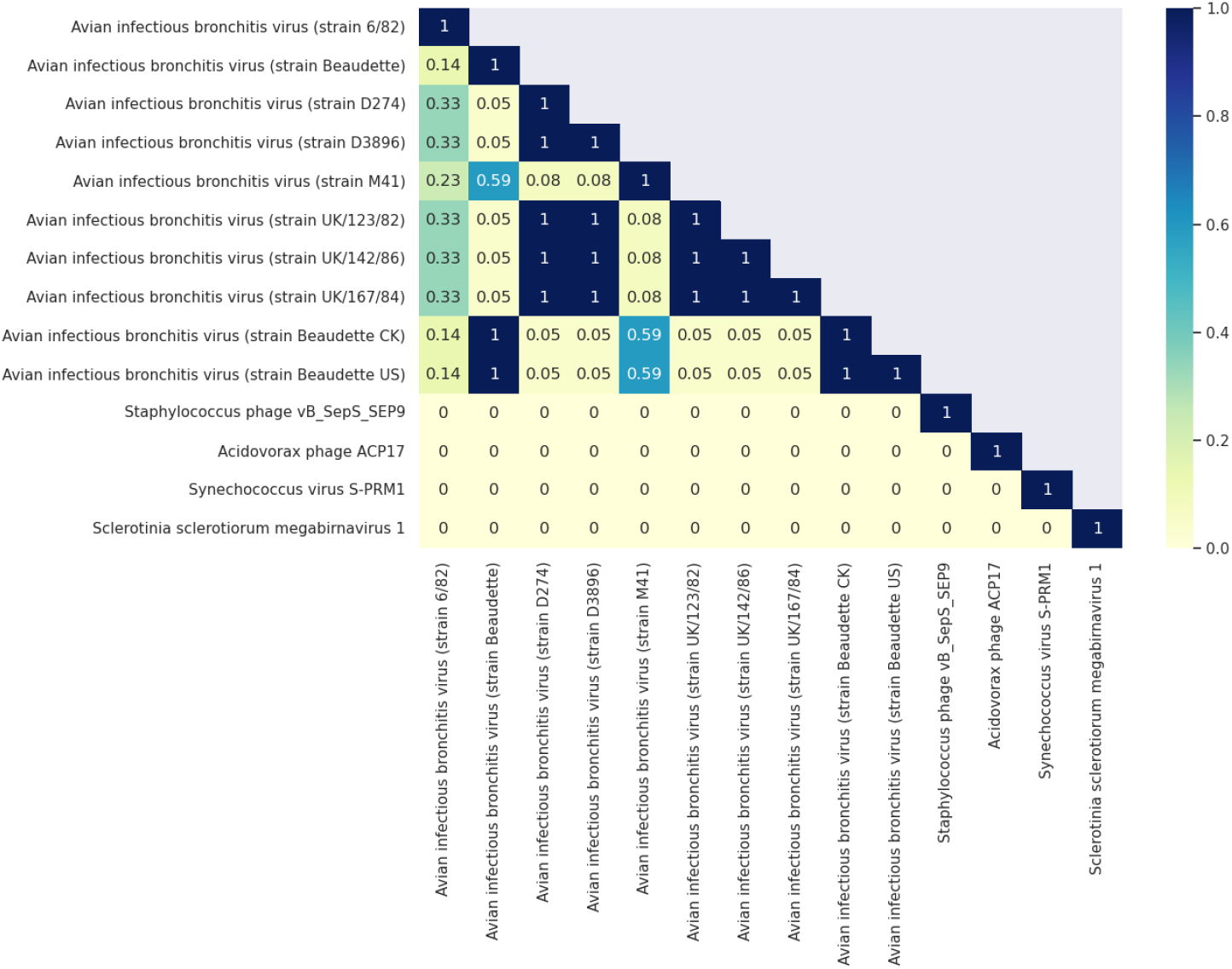
Detected peptidome similarity of the top scoring viral strains for the avian bronchitis sample. Peptides included for the peptidome similarity computation are restricted to the peptides that were identified by SearchGUI/PeptideShaker and included in the graphical model.

For the herpesvirus, PepGM wrongly identifies human herpesvirus strain 17 as present. The detailed scored results can be seen in the supplementary material in Figure 7. The second and third highest scoring taxa, while also human herpesvirus 1 strains, are the neighbouring strains RH2 and KOS. The correct herpesvirus strain, strain F, appears in 4th position with a low score of 0.15. Using TaxIt, this human herpesvirus was misidentified as well, and the authors suspected that the sample annotation might be incorrect. As the strain used is clearly described in the associated publication, ^41^ we believe this to be unlikely. The human herpesvirus strain identified, strain 17, is the most commonly used reference strain for human herpesviruses and is therefore strongly over-represented (2553 entries in NCBI Protein) in the general NCBI reference database compared to strain F (only 282 entries in NCBI Protein). Again, the score PepGM attributes to its identification, about 0.81, is lower than the lowest score of the correct identifications, highlighting once more the usefulness of the attributed scores.

To investigate the effect of missing species information on PepGM, we evaluated cowpox virus PXD014913, the adenovirus, and the hendravirus using the RefSeq Viral database where the entries corresponding to the correct species and strain (if present) have been removed. Plots of the results can be found in the supplementary materials in Figure 15. For the cowpox virus, a range of Orthopoxviruses is identified with confidence scores around 0.6, signalling poor results. Similar results - a series of low confidence assignments of strains from the species human mastadenovirus D (whereas the correct species would have been human mastadenovirus C) can be observed for the human adenovirus 2, when the correct species (human mastadenovirus C) has been removed from the database. A series of low confidence identifications from the same genus therefore indicates that a microorganism of that genus is present, but that the correct species is unknown or missing in the database. For the hendravirus however, the species Nipah Henipavirus, from the correct genus Henipavirus, is identified with a confidence score of 0.86 (the correct strain - Hendra Henipavirus, was assigned a score of 0.99). In this case, the PepGM score does not give a clear indication that the identification might be erroneous.

### Effect of unique or shared peptides and their score on PepGM results

To investigate how the number of unique or shared peptides, and thus the peptidome similarity, and their confidence score per taxon affect our results, we run PepGM using the graph structure resulting from the avian bronchitis sample with different, manually added peptides. The avian bronchitis graph structure is most suited for this investigation as all three possible strains have the same detected peptidome, and thus are not differentiable in the graph structure. The manually added peptide constellations are:

1. A single high confidence peptide node (score 0.9) added to the Beaudette US strain
2. A single medium confidence peptide node (score 0.7) added to the Beaudette US strain
3. A single lower confidence peptide node (score 0.5) added to the Beaudette US strain
4. A shared medium-confidence peptide node (score of 0.7) added to the Beaudette CK and US strains

Table 2 shows how these added peptides affect the results. Adding a a single high confidence peptide for the avian bronchitis Beaudette US strain elevates its score to 0.92, and lowers the score attributed to its two neighboring strains to 0.62, demonstrating that a single high confidence peptide is sufficient for PepGM to recognise the correct strain, without completely discarding the two other possible strains. Adding a medium confidence peptide does not increase the score of 0.78 for the Beaudette US strain, but lowers the scores attributed to the other strains to 0.58, meaning PepGM is still able to recognize the present strain. Adding a peptide that is equally as likely to be correct as it is to be false does not help detection of the present viral strain, and finally, adding a medium-confidence peptide shared between two neighboring strains increases their attributed score to 0.8 while reducing the score attribute to the third neighbor strain. The results show that PepGM sensitively takes into account peptide scores, and, depending on the confidence of the detected peptides, a single unique peptide is sufficient to be able to distinguish strains with otherwise equal detected peptidomes.

**Table 2:** Variations in the scores of the three best scoring strains for the avian bronchitis sample for different added peptide constellations

| Added peptide constellations |  | 1 | 2 | 3 | 4 |
| --- | --- | --- | --- | --- | --- |
| 3 top scoring strains & their score | Beaudette US | 0.92 | 0.78 | 0.78 | 0.8 |
|  | Beaudette CK | 0.62 | 0.58 | 0.78 | 0.8 |
|  | Beaudette | 0.62 | 0.58 | 0.78 | 0.61 |

We also investigated, for the herpesvirus, the avian bronchitis, and the hendravirus, the effect of setting different FDR thresholds-1%, 5% and 20%. These effects were very limited, as shown in the supplementary materials (Figure 16). As described for manually added peptides, the taxonomic identifications and scores computed by PepGM are dominated by high-confidence peptides, which explains this insensitivity to changes in FDR threshold.

### Effect of optional host filtering

After demonstrating the usefulness of PepGM, we investigate the effect of adding the host filtering step (described as step 0 above). In some cases, such as for both SARS-CoV-2 samples shown in Section S2, Figures 9 to 12, adding the host filtering step does not or only slightly alter the taxonomic identification results. In other cases, the host filtering step does alter the identification results. For the human adenovirus, a comparison of Figure 2 (without host filtering) and Figure 1 (with host filtering) in Section S2 of the supplementary material shows that the host filtering decreases the identification accuracy - for the filtered spectra, PepGM is not able to differentiate between the human adenovirus strains 2, 6 and 5 anymore. It is likely that filtering for possible host and contaminant peptides removed some peptides that also mapped to one of either strains. A similar effect can be seen in Figure 2 in Section S2 of the supplementary material where the filtering reduces the ability of PepGM to differentiate between different avian bronchitis strains. For purified viral samples, as is the case for all samples analyzed in this benchmark, we therefore do not recommend to use the host filtering step.

### Graphical model parameter evaluation

We also investigate the parameters *α, β* and *γ* that were identified as best using the metric described in Section Methods. For each sample, the chosen parameters are shown in Table 3. For the parameter *γ*, representing the prior probability for a taxon to be present, the grid search was conducted in the range *γ* ∈ {0.1, 0.3, 0.5}. Here, setting *γ* = 0.5, which corresponds to a uniform distribution, encodes the knowledge that no prior information is available on any taxon. Table 3, however, shows that for most samples *γ* = 0.1 is the selected parameter. This is a reflection of the fact that we are dealing with single organism samples where indeed, for most taxa, the true prior would be close to 0 as they are absent. For the avian bronchitis, the grid search identified *γ* = 0.3 as the best fitting prior, which, in this case, is caused by the simultaneous ‘presence’ of the three Beaudette avian bronchitis strains between which all peptides are shared, and PepGM thus cannot differentiate.

**Table 3:** Parameters of the noisy OR conditional probability tables, *α, β* and *γ*, selected as best fitting for each sample.

| Sample | Virus (strain) | $\alpha$ | $\beta$ | $\gamma$ |
| --- | --- | --- | --- | --- |
| PXD003013 | Cowpox virus (Brighton Red) | 0.01 | 0.01 | 0.1 |
| PXD014913 | Cowpox virus (Brighton Red) | 0.05 | 0.01 | 0.1 |
| PXD018594 | SARS-CoV-2 | 0.6 | 0.7 | 0.1 |
| PXD025131 | SARS-CoV-2 | 0.4 | 0.7 | 0.1 |
| PXD005104 | Human herpesvirus 1 (strain F) | 0.01 | 0.4 | 0.1 |
| PXD002936 | Avian bronchitis (Beaudette CK) | 0.05 | 0.1 | 0.3 |
| PXD001165 | Hendra virus 1994 | 0.05 | 0.05 | 0.1 |
| PXD004095 | Human adenovirus (strain 2) | 0.01 | 0.1 | 0.1 |

For the parameters *α* (corresponding to the ‘emission probability’ of a peptide given its parent taxons presence) and *β* (the false detection probability of a peptide), the observed ranges, namely *α* ∈ [0.01, 0.4], *β* ∈ [0.01, 0.7] and *α* ≤ *β* are somewhat harder to explain. If interpreted according to their meaning in the noisy OR model, the parameters would suggest that the probability of observing a peptide, given the presence of its parent taxon, is very low, and that many peptides are detected erroneously. However, we modeled a complex physical process with a simple noisy OR approach and made the assumption that *α* and *β* are the same for all peptides, both of which are strong simplifications. It is therefore more likely that our simplified noisy OR model is not an accurate representation of the underlying peptide generation process, which impairs the direct interpretation of the *α* and *β* values. Nevertheless, given the effectiveness of PepGMs taxonomic identification, the use of this simplified model remains justified.

### Comparison to the identification results of TaxIt

We previously mentioned several other taxonomic identification pipelines that can achieve strain-level resolution. Of these, TaxIt is the only one able to reliably provide strain-level identifications for a broad range of samples.^19^ We therefore compared the identification performance, runtime and memory use of PepGM and TaxIt.

Table 4 shows that PepGM identified the viral train equally as well as TaxIt for 6 out of 8 viral samples. In the two cases where PepGMs identifications were ambiguous (the avian bronchitis sample) or correct regarding the species, but incorrect regarding the strain (the herpesvirus sample), the identifications made by TaxIt were incorrect or ambiguous as well. The issues of TaxIt with correctly identifying the taxonomy of these two samples likely have the same origins as those affecting PepGM: for the avian bronchitis, the detected peptidomes were too similar; for the herpesvirus, the incorrect strain was overrepresented in the reference databases.

**Table 4:**
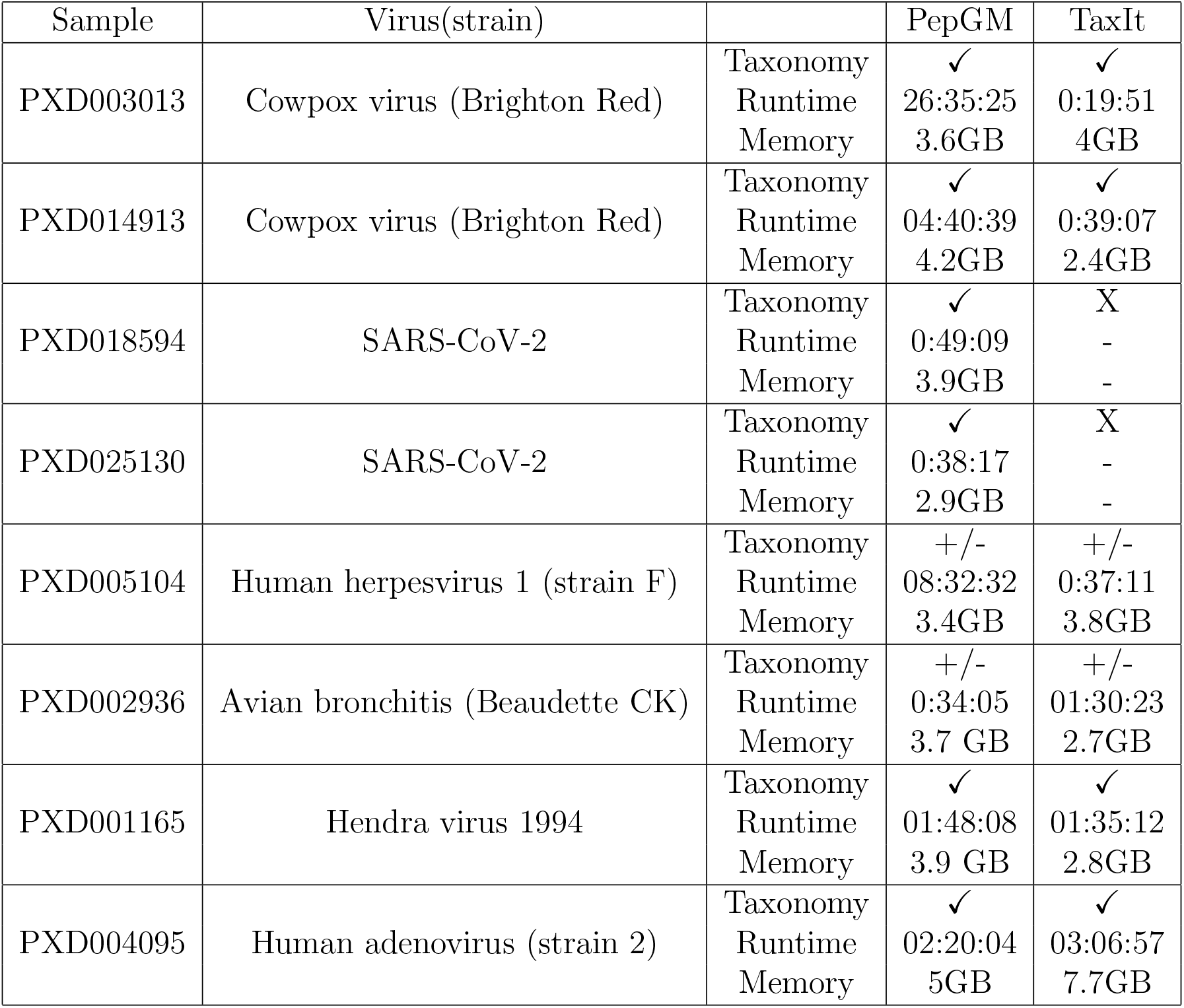
Comparison of identification results made by PepGM and TaxIt, runtime and memory use. ✓ means correct identification, +/− means correct species, but incorect strain or ambiguous strain attribution, and X means the analysis failed.

Notably, PepGM is able to achieve a similar taxonomic resolution using only the peptide search results from a single database search, while TaxIt relies on a second, refined search. PepGM has the added benefit of providing confidence scores, indicating when its identifications might be less accurate. For the two coronavirus samples, TaxIt did not produce results - this is due to its use of the NCBI Entrez API and the currently not updated NCBI taxonomy structure for certain taxa in the NCBI nodes dump files. TaxIt identified the Severe Acute Respiratory Syndrome-related coronavirus as present species and subsequently tried to download all corresponding strain level proteins through the API. However, the 260 corresponding strains were too much to be queried through the API. PepGM, using only the NCBI protaccession2taxid file to identify potentially present species, directly recognized SARS-CoV-2 as a potential species and therefore queried much less information through the API, allowing it to finish the analysis.

The total runtime and memory usage of PepGM and TaxIt are shown in Table 4. The memory use of PepGM and TaxIt was comparable. Regarding runtime, we can not make one statement for all samples. In our benchmark, TaxIts runtime was often dominated by the time necessary to download individual protein entries through the NCBI API. PepGM, in contrast, queries whole proteomes per organism at once. Therefore, PepGMs runtime is more dependent on the number of strains and peptides included in the graph. Building the factor graph requires querying pairwise peptide-taxon proteome comparisons and the more complex statistical computations, where messages are updated iteratively for each graph edge, slow down as the graph size increases. PepGMs grid search can in theory be executed completely in parallel and could thus be accelerated markedly by using more CPU cores. Should this not be possible, one could choose to reduce the number of points searched through in the grid search, possibly affecting results accuracy but leading to an important speed-up. For two samples, cowpox virus PXD003013 and the cowpox virus sample, the runtime was a lot longer. For both, it was dominated by few belief propagation runs on certain parameter sets that took a very long time to converge. In fact, some initial variable sets can cause oscillations in the messages passed through belief propagation, slowing down and possibly impeding convergence.^42^ Adding some form of optional damping^43^ could, in the future, speed up the belief propagation algorithm for the variable sets in question.

## Discussion and Outlook

We have shown that PepGM consistently and reliably determines the taxonomy of viral samples with strain-level accuracy. Using a graphical model approach, PepGM makes use of peptide scores returned by database search engines or post-processing tools to calculate meaningful confidence scores for its taxonomic assignments. This could be especially useful in clinical settings, when knowing the reliability of provided strain assignments could influence therapeutic decision making.

For two of the viral samples, the taxonomic assignments were either ambiguous or identified the correct species, but only a closely related viral strain. For the avian bronchitis, PepGM was not able to differentiate between the three Beaudette subtypes. We showed that this was due to their detected peptidomes being identical. Because PepGM indicated the lower confidence of its assignment by a lower taxon score, this remains a defensible result, especially as TaxIt was not able to resolve the ambiguity either. For the herpesvirus, PepGM identified strain 17 instead of strain F as present in the sample. No other algorithm was able to provide a correct taxonomic assignment for this sample,^19^ and neither was TaxIt in our own benchmark. As previously discussed, the misassignment can be explained by the over representation of strain 17 in the available proteome references.

More generally, PepGM does not overcome the reliance on public reference databases. By combining the results of a search against the curated reference RefSeq Viral with the query of all available strain proteomes to potentially include in the peptide-taxon graph, PepGM aims to optimally use the available information. Uncurated references, which we include by querying all available strain-level reference proteomes, can be error prone.^44^ Our graphical model approach, which takes into account the probability of false peptide identifications (and assignment to taxa), is able to partially mitigate this issue.

When the correct species was removed from the reference database, PepGM assigned various species from the correct genus with low confidence scores, except for the hendravirus, where the Nipah Henipavirus was incorrectly assigned - still with a slightly lower probability. This shows again that the score provided by PepGM indicates the correctness of a taxonomic assignment well, but also that assigning a threshold score above which PepGM is certain of its assignment is futile. More generally, many taxonomic assignments with similar, low scores, seem to indicate that the correct strain or species was not in the database used, while a few high, but close-scoring taxonomic assignments would indicate that it was not possible to resolve between neighbouring strains (as was the case for the avian bronchitis sample).

Closer examination of the graphical model parameters, identified from the grid search results, revealed that the value taken by the parameters *α* and *β* seem abstract related to their meaning in the noisy OR model. Future development could aim at using peptide-specific *α* and *β* parameters based, for example, on the prediction of peptide probability after tryptic cleavage^45^ for *α* or posterior error probabilities, derived from the PSMs but aggregated at peptide level, for the parameter *β*.

Compared to other available tools that can be used for strain-level identification of viral proteome samples, PepGM reaches at least the taxonomic resolution of TaxIt, the previously best performing approach, with the additional benefit of providing confidence scores. Memory wise, PepGM and TaxIt are comparable, but for the runtime, there are sample specific differences. For PepGM, as described previously, both the construction of the factor graph and the belief propagation runtimes increase with the graph size - meaning the more peptides and taxa have to be included in the graph, the slower PepGM. This is a disadvantage compared to TaxIt - which only becomes very slow when an excessive number of taxa have to be queried through the NCBI API. The runtime of PepGM, except for two samples, did not exceed 4 hours. This can still be a hindrance in clinical settings, were fast decisions need to be made. A speed-up could be achieved in the future by improving the belief propagation algorithm. As mentioned previously, certain initial variable sets can cause oscillations slowing down and possibly impeding convergence. Damping these messages could be a future solution, as was done for the protein inference algorithm based on belief propagation. ^29^

For now, PepGM is limited to the taxonomic identification of viral samples, but the graphical model approach should be extensible to all types of organisms. Again, in the clinical context, the strain-level identification of pathogenic bacterial samples is of particular interest. To make this possible, PepGM could be extended by using a general bacterial reference database, such as RefSeq for bacteria, and including the peptide-taxon mapping information for bacteria as well. Because strain-level peptides have to be downloaded through the NCBI API and because PepGM downloads the strain-level information for all target species included in the graph, it would likely be necessary to set up local database options to optimize speed and circumvent the NCBI API. Another alternative would be to combine PepGM with the query of pre-digested peptide databases such as Unipept, where tryptic peptides are pre-annotated taxonomically. Another potential area of application would be metaproteomics, where complex microbial mixtures have to be taxonomically and functionally identified. Peptide-taxon and peptide-function relationships can be represented in bipartite graphs, and a similar graphical model could be applied. Yet, in such a scenario, the metric used to identify the best graphical model parameters would need to be adapted to suit the presence of multiple organisms at once.

Finally, the application of PepGM for the strain-level identification of viral proteomes faces the issue, in practice, of the small size of viruses and therefore the low abundance of viral proteins.

## Conclusion

In this work, we have presented PepGM, a new graphical model based workflow for the strain-level taxonomic assignment of viral proteome samples. We have shown that PepGM is able to consistently provide accurate taxonomic identifications with associated, meaningful confidence scores, a feature that could be especially helpful in the clinical context, when medical decisions have to be made based on the confidence of the identification results. More generally, we have demonstrated the usability of graphical models for the taxonomic identification of proteomes samples using the restricted example of viral samples.

## Acknowledgements

TH was supported by the German Research Foundation (DFG) (MU 4430/2-1) and acknowledges additional support by the Joachim-Herz-Foundation. TH and TM gratefully acknowledge support and helpful comments by their colleagues Yannek Nowatzky and Kristin Vogel. LM acknowledges the Research Foundation - Flanders (FWO) [G028821N to L.M.], the European Union’s Horizon 2020 Program (H2020-INFRAIA-2018-1) [823839 to L.M.], and Ghent University Concerted Research Action [BOF21-GOA-033 to L.M.]

## Notes

### Competing Interest Statement

The authors have declared no competing interest.

