## Supplementary Material for "PepGM: A probabilistic graphical model for taxonomic inference of viral proteome samples with associated confidence scores"

†*s.3 eScience, Bundesanstalt für Materialforschung und - Prüfung, Berlin, Germany*

‡*VIB-UGent center for Medical Biotechnology, VIB, Belgium*

¶*Department of Biomolecular Medicine, Ghent University, Belgium*

**S3 Additional tables and figures** **5**

Table 1: Packages used in the PepGM workflow

| Package name | reference |
|---|---|
| numba | [1] |
| Ete3 | [2] |
| mmh3 | [3] |
| pandas | [4] |
| numpy | [5] |
| networkx | [6] |
| scipy | [7] |
| Biopython | [8] |
| matplotlib | [9] |
| seaborn | [10] |

### S1 Libraries used for PepGM

### S2 Search parameters for the publicly available samples

#### Human adenovirus 2 sample

The human adenovirus 2 sample was downloaded from PRIDE (PXD004095) and the sample 20140812_LC1_Sara_24h_MH+AL_10-03.raw was selected for analysis. We converted the .raw file to an .mgf file using the ProteoWizard MSConvertGUI (version 3.0.21232). In accordance with the original processing protocol, for the SearchGUI parameters, we used a tryptic search, 2 allowed missed cleavages, a fragment ion mass tolerance of 20ppm and precursor ion mass tolerance of 4.5ppm. For the modifications, we used the fixed modification cysteine carbamidomethylation (+57 Da) as well as additional variable modifications methionine oxidation (+16 Da), acetylation of the protein N-terminus (+42 Da) and phosphorylation of serine, threonine and tyrosine (+80 Da). The host proteome - homo sapiens, was downloaded from Uniprot and had 26591 entries.

#### Avian bronchitis (Beaudette CK) sample

The avian bronchitis (Beaudette CK) sample was downloaded from PIRDE (PXD002936) and we used the sample BeauR2.raw for analysis. We converted the .raw file to an .mgf file using the ProteoWizard MSConvertGUI (version 3.0.21232). For the SearchGUI parameters, we used a tryptic serach, 2 allowed miscleavages, a precursor ion tolerance of 100ppm, a fragment ion tolerance of 0.5Da and a charge of 2-6 )in accordance with the default settings of X!Tandem). As fixed modification, we specified cysteine carbamidomethylation (+57 Da). The host proteome, chicken or gallus gallus, was downloaded from Uniprot and had 2618 entries.

#### Human herpesvirus 1 strain F

The human herpesvirus 1 (strain F) sample was downloaded from Pride (PXD005014) and we selected the sample OR10_20151030_EC_HHV1_A_02.raw for analysis. We converted the .raw file to an .mgf file using the ProteoWizard MSConvertGUI (version 3.0.21232). For the SearchGUI parameters, in accordance with the original processing parameters, we used a precursor ion tolerance of 100ppm, a fragment ion tolerance of 0.5Da and a charge of 2-6 (in accordance with the default settings of X!Tandem). As fixed modification, we specified cysteine carbamidomethylation (+57 Da) and variable modification methionine oxidation (+16 Da). The host proteome - homo sapiens, was downloaded from Uniprot and had 26591 entries.

#### Hendra virus sample

Hendra henipavirus samples of the strain horse/Australia/Hendra/1994 were downloaded from Pride (PXD001165) and we selected the sample Slice_10_human.raw for analysis. We converted the .raw file to an .mgf file using the ProteoWizard MSConvertGUI (version 3.0.21232). For the SearchGUI parameters we used a precursor ion tolerance of 100ppm, a fragment ion tolerance of 4.5ppm and a charge of 2-6 (in accordance with the default settings of X!Tandem). As fixed modification, we specified cysteine carbamidomethylation (+57 Da). As the sample is SILAC labeled we additionally specified Arginine 13C(6) 15N(4), and Lysine 13C(6) 15N(2) as variable modifications. The host proteome - homo sapiens, was downloaded from Uniprot and had 26591 entries.

#### Cowpox virus strain Brighton Red

We downloaded two Cowpox Virus (strain Brighton Red) samples from Pride.
The first one is the sample PXD003013, where we selected the spectrum CPXV_BR_2.RAW for analysis. We converted the .raw file to an .mgf file using the ProteoWizard MSConvertGUI (version 3.0.21232). In accordance with the original processing protocol, for the SearchGUI parameters we used a tryptic search, a precursor ion tolerance of 4.5ppm, a fragment ion tolerance of 0.5Da. As fixed modification, we specified cysteine carbamidomethylation (+57 Da) as well as additional variable modifications methionine oxidation (+16 Da), acetylation of the protein N-terminus (+42 Da). The host proteome - homo sapiens, was downloaded from Uniprot and had 26591 entries.
The second is from the Pride project PXD014913, where we selected the sample CPXV-0,1MOI-supernatant-HEp-24h.mgf for analysis. We converted the .raw file to an .mgf file using the ProteoWizard MSConvertGUI (version 3.0.21232). In accordance with the original processing protocol. For the SearchGUI parameters we used a tryptic search, a precursor ion tolerance of 4.5ppm, a fragment ion tolerance of 0.5Da. As fixed modification, we specified cysteine carbamidomethylation (+57 Da) as well as additional variable modifications methionine oxidation (+16 Da). The host proteome - homo sapiens, was downloaded from Uniprot and had 26591 entries.

#### SARS-CoV-2

The first SARSs-CoV-2 sample was downloaded from Pride (PXD018594) ad we selected the sample Q10467_MS20-17_CoV2_J4(MOI-01)r.mgf for analysis. According to te original processing protocol, we used a tryptic search, a precursor ion tolerance of 4.5ppm, a fragment ion tolerance of 0.5Da. As fixed modification, we specified cysteine carbamidomethylation (+57 Da) as well as additional variable modifications methionine oxidation (+16 Da).
We downloaded the second SARS-CoV-2 sample from Pride (PXD025130) and selected the sample Q26431_MS20-025_Virus-purif_1.mgf for analysis. According to the original processing protocol, we used a tryptic search, a precursor ion tolerance of 4.5ppm, a fragment ion tolerance of 0.5Da. As fixed modification, we specified cysteine carbamidomethylation (+57 Da) as well as additional variable modifications methionine oxidation (+16 Da)

### S3 Additional tables and figures

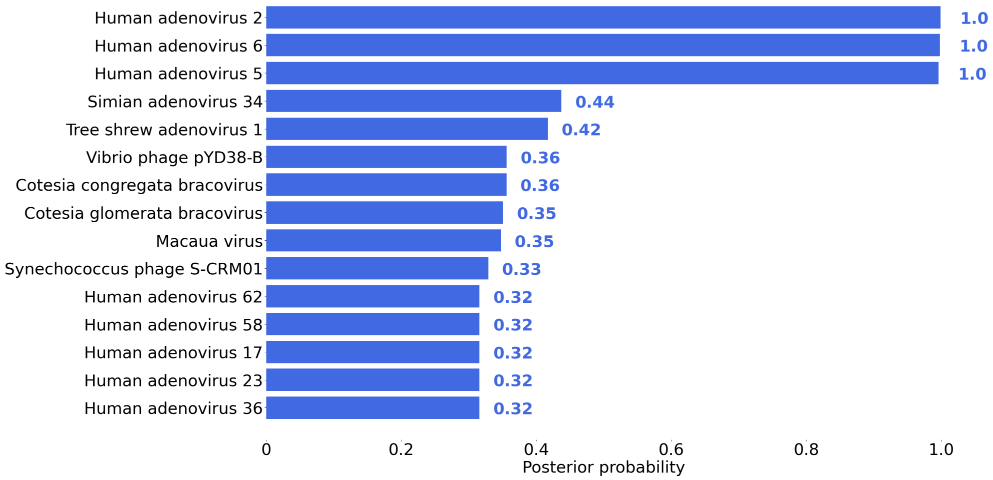

Figure 1: PepGM identification Results for the adenovirus sample with additional host filtering. Bar plot representation of the 15 highest scoring taxa, with the rounded attributed score written next to each bar.

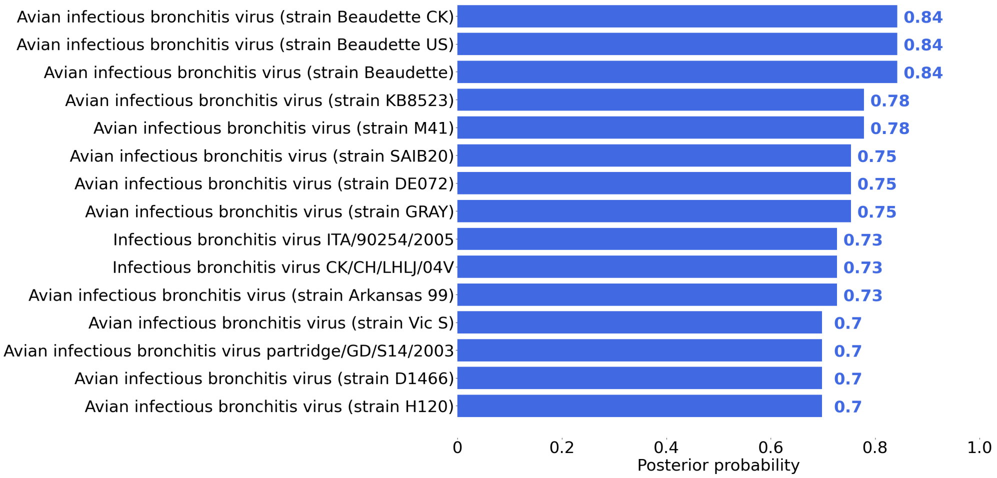

Figure 2: PepGM identification Results for the avian bronchitis sample with additional host filtering. Bar plot representation of the 15 highest scoring taxa, with the rounded attributed score written next to each bar.

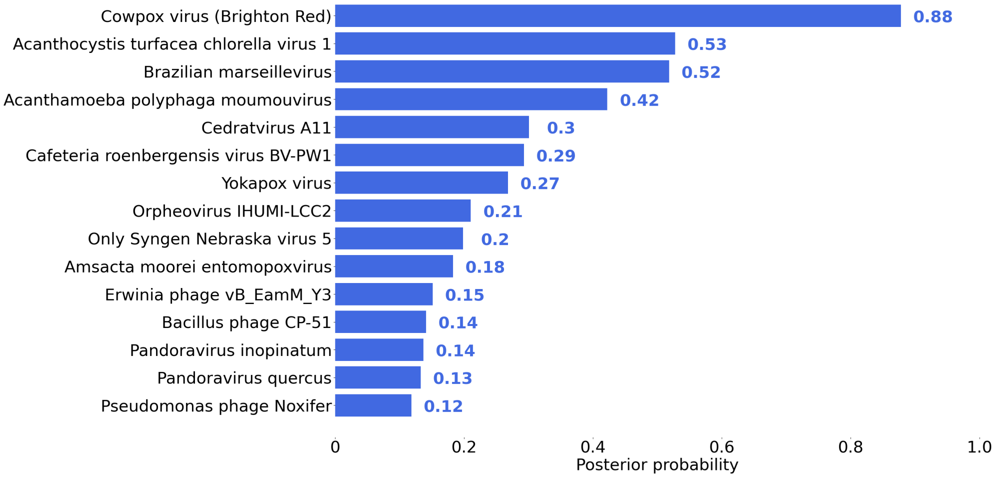

Figure 3: PepGM identification Results for the cowpox sample PXD014913 Bar plot representation of the 15 highest scoring taxa, with the rounded attributed score written next to each bar.

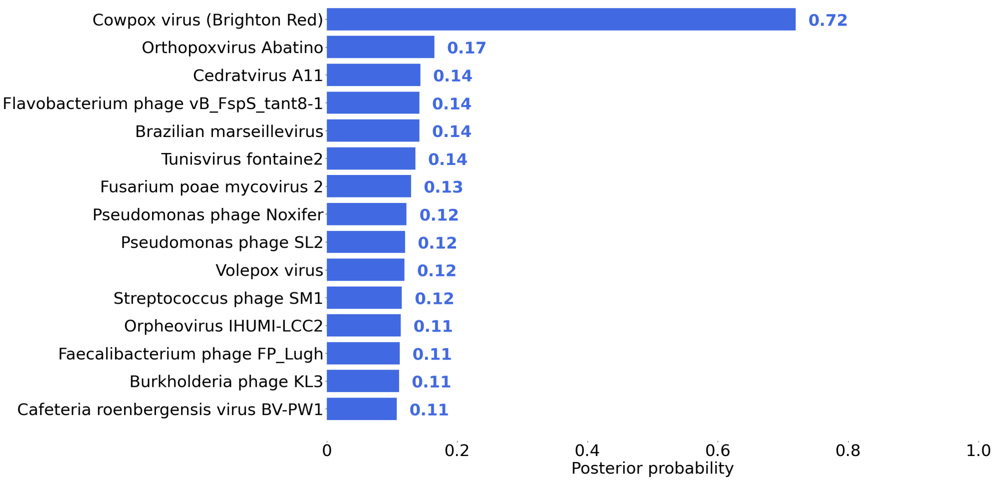

Figure 4: PepGM identification Results for the cowpox sample PXD014913 with additional host filtering. Bar plot representation of the 15 highest scoring taxa, with the rounded attributed score written next to each bar.

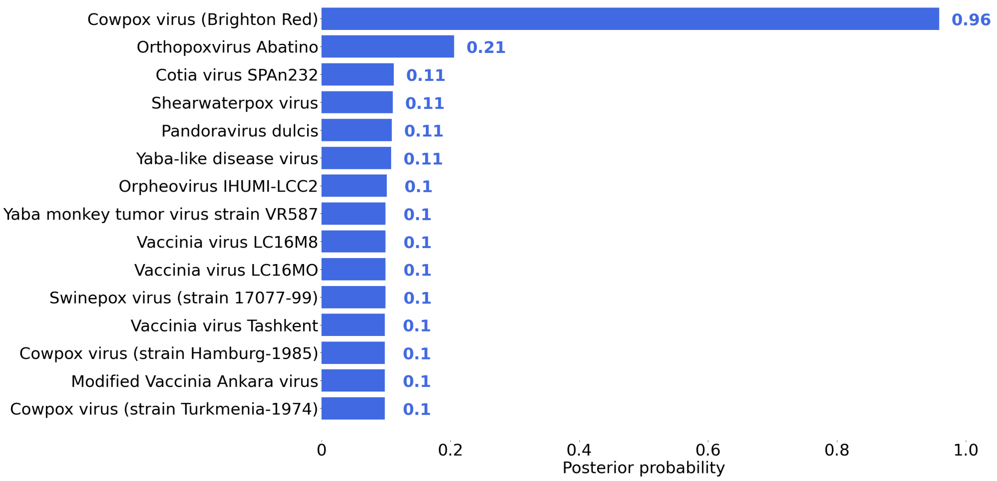

Figure 5: PepGM identification Results for the cowpox sample PXD003013 Bar plot representation of the 15 highest scoring taxa, with the rounded attributed score written next to each bar.

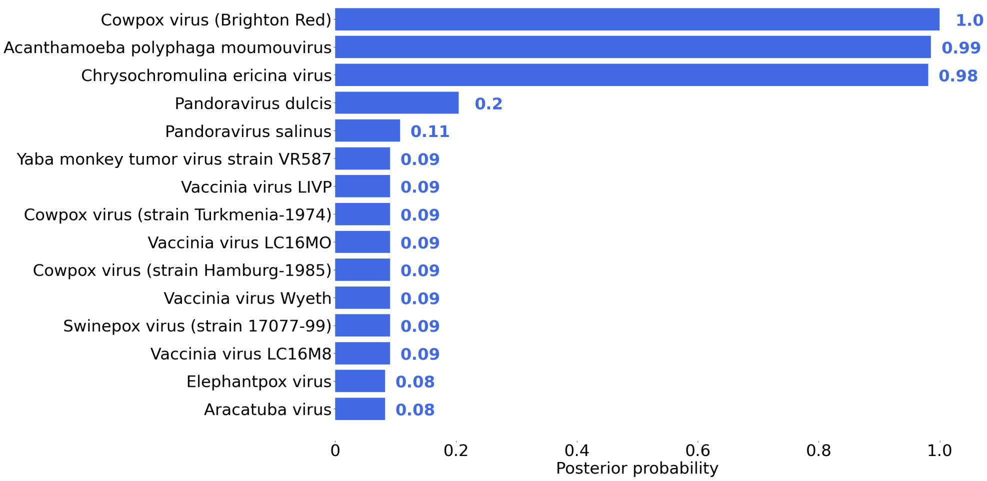

Figure 6: PepGM identification Results for the cowpox sample PXD003013 Bar plot representation of the 15 highest scoring taxa, with the rounded attributed score written next to each bar.

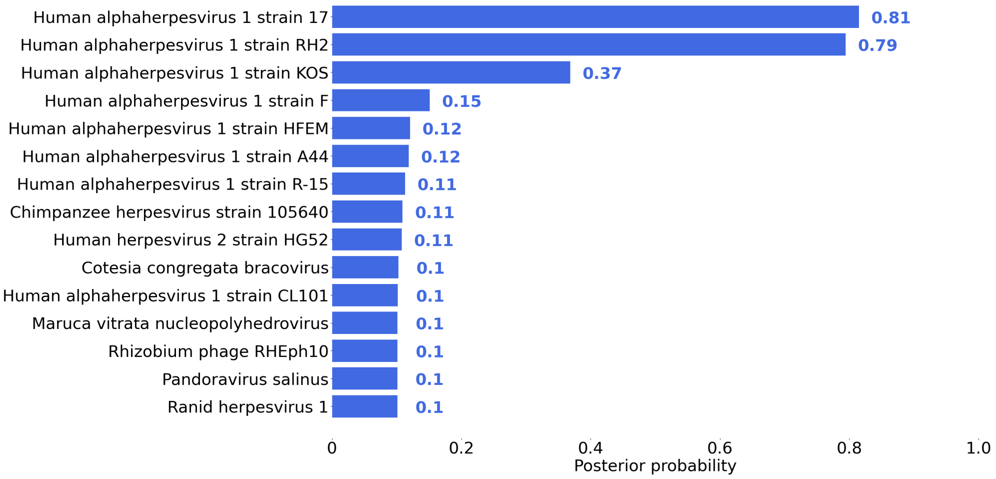

Figure 7: PepGM identification Results for the human herpesvirus sample. Bar plot representation of the 15 highest scoring taxa, with the rounded attributed score written next to each bar.

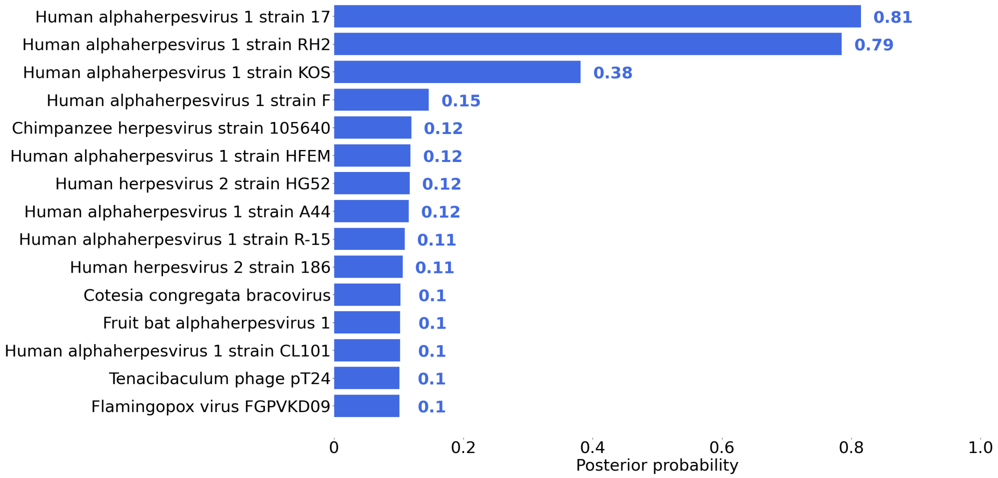

Figure 8: PepGM identification Results for the human herpesvirus sample with additional host filtering. Bar plot representation of the 15 highest scoring taxa, with the rounded attributed score written next to each bar.

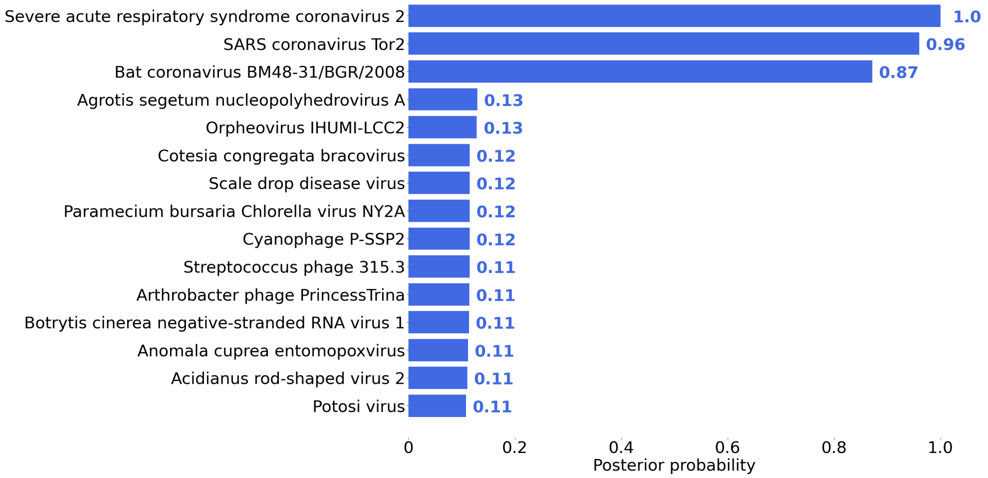

Figure 9: PepGM identification Results for the SARS-CoV-2 sample PXD025130. Bar plot representation of the 15 highest scoring taxa, with the rounded attributed score written next to each bar.

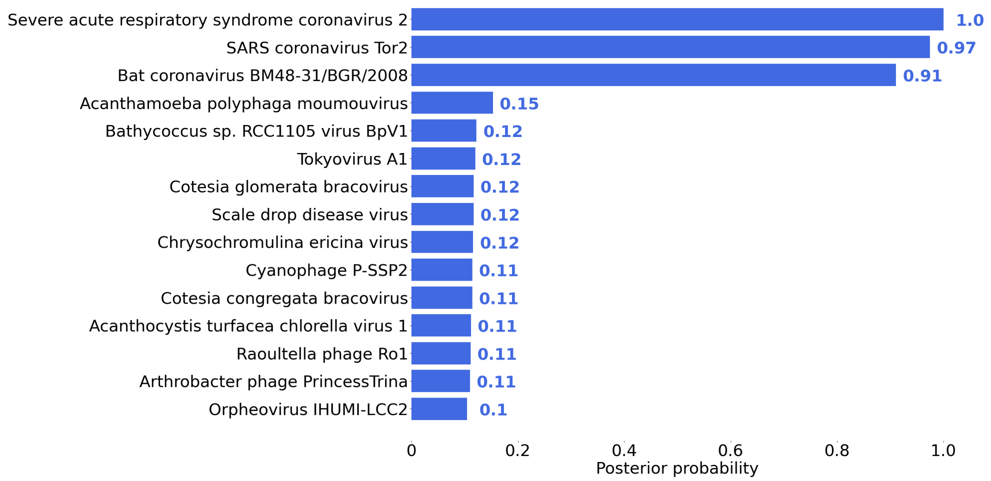

Figure 10: PepGM identification Results for the SARS-CoV-2 sample PXD025130 with additional host filtering. Bar plot representation of the 15 highest scoring taxa, with the rounded attributed score written next to each bar.

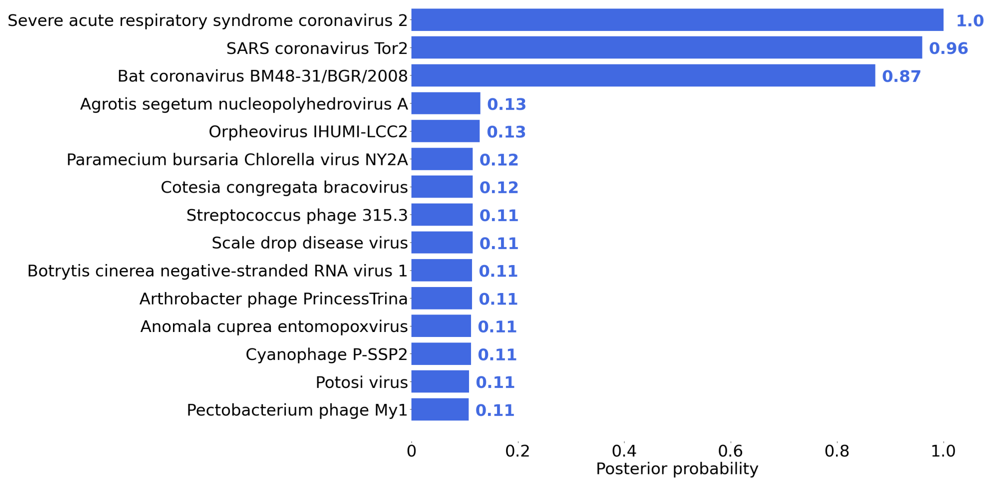

Figure 11: PepGM identification Results for the SARS-CoV-2 sample PXD025131 sample. Bar plot representation of the 15 highest scoring taxa, with the rounded attributed score written next to each bar.

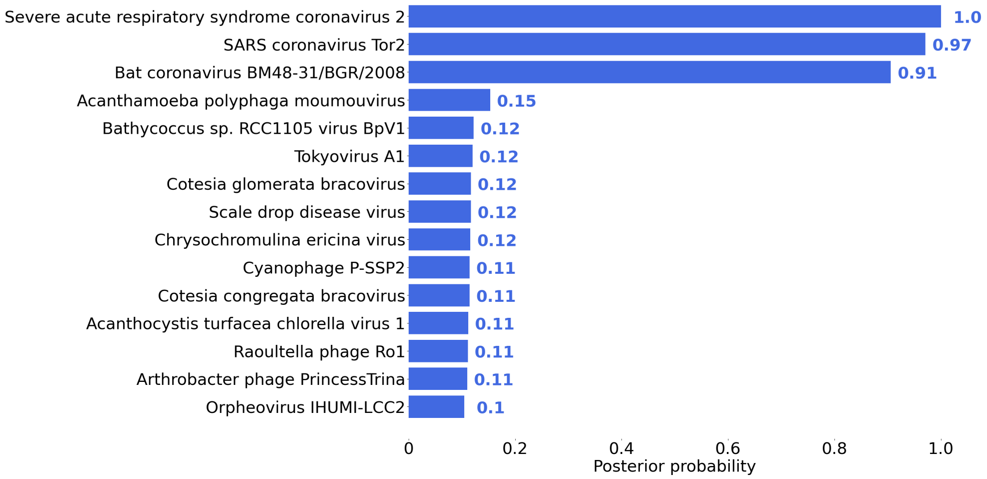

Figure 12: PepGM identification Results for the SARS-CoV-2 sample PXD025131 sample with additional host filtering. Bar plot representation of the 15 highest scoring taxa, with the rounded attributed score written next to each bar.

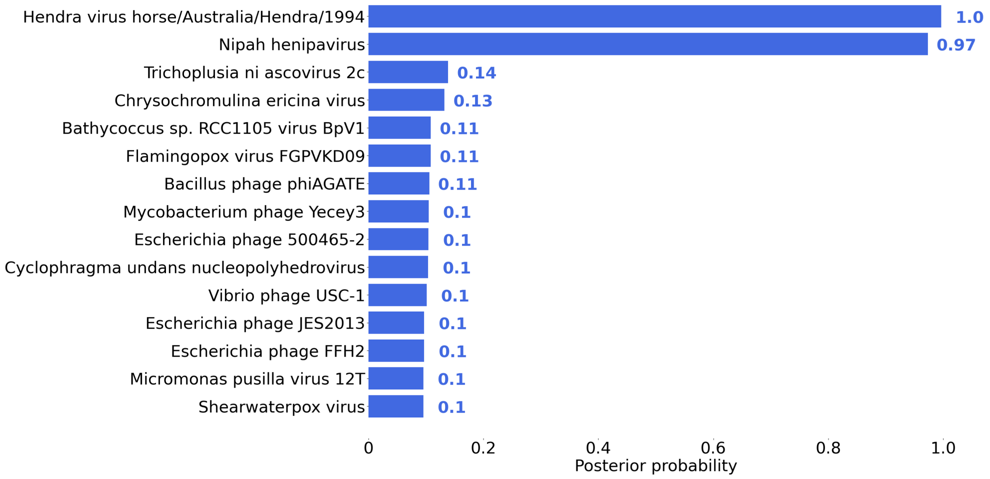

Figure 13: PepGM identification Results for the hendravirus sample. Bar plot representation of the 15 highest scoring taxa, with the rounded attributed score written next to each bar.

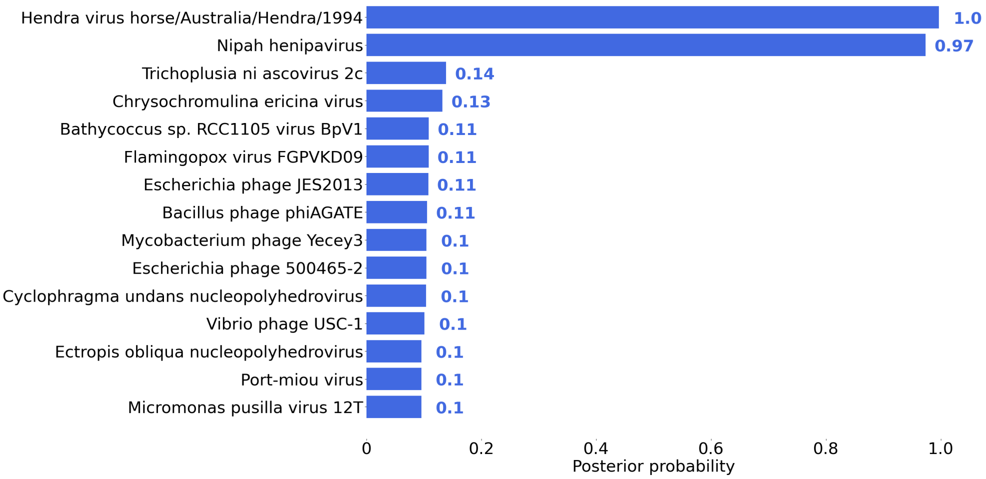

Figure 14: PepGM identification Results for the hendravirus sample with additional host filtering. Bar plot representation of the 15 highest scoring taxa, with the rounded attributed score written next to each bar.

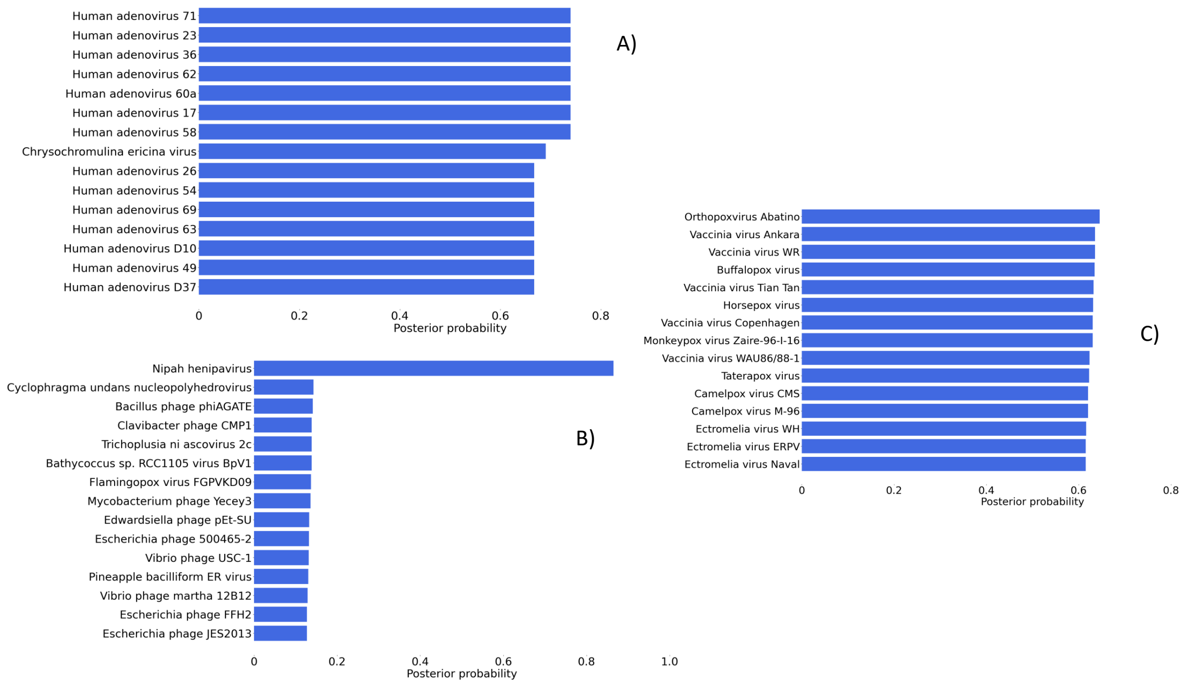

Figure 15: PepGM identification Results for A) the adenovirus sample, B) The hendravirus sample and C) the cowpox sample. For each, the correct species and/or strain (if present in the the RefSeq Viral database) was removed.

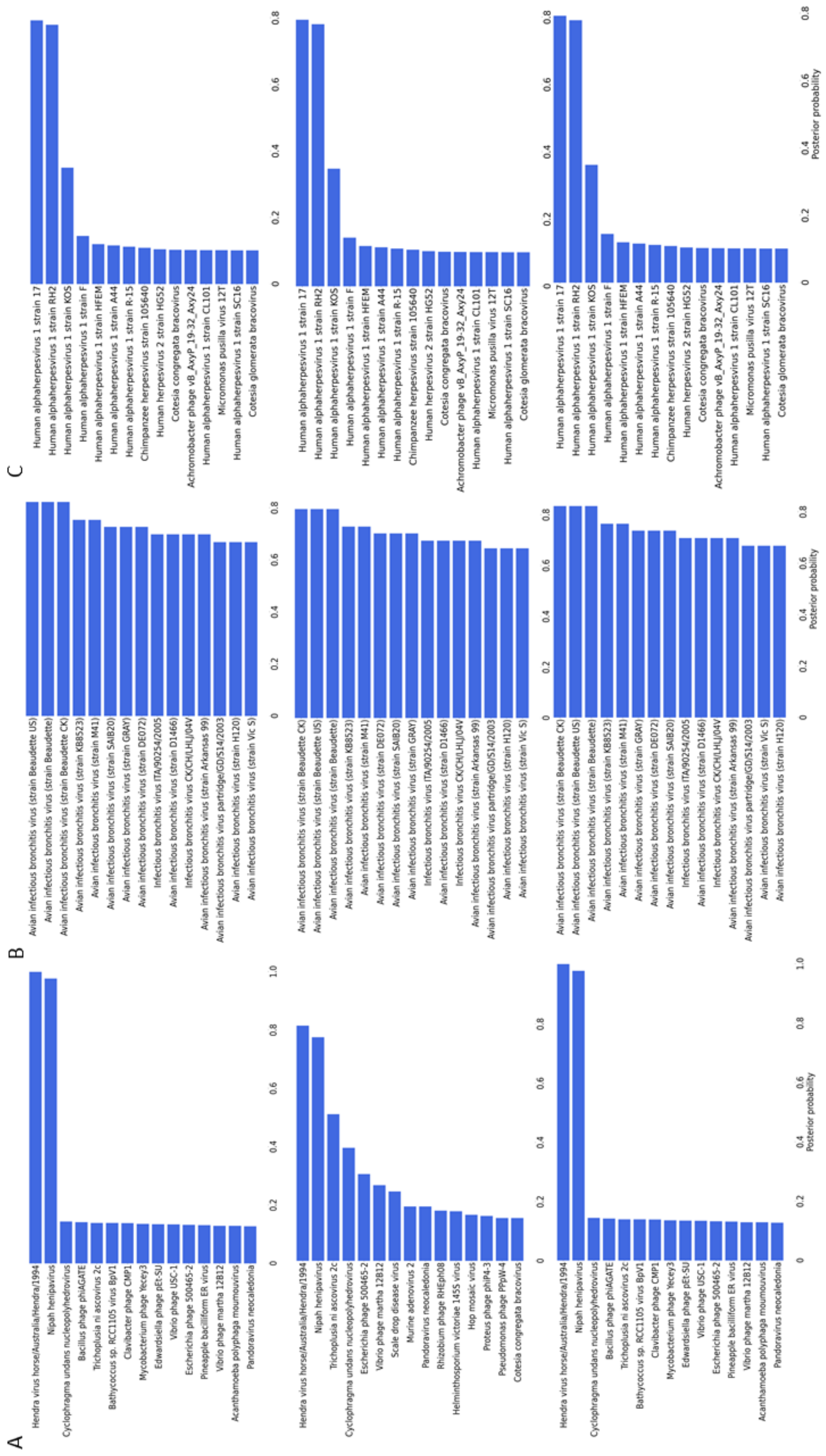

Figure 16: 2.4 PepGM results for (A) the hendravirus, (B) the avian bronchitis and (C) the herpesvirus sample. FDRs of 1%, 5% and 20% (top to bottom panels) were used for the database search step.
